# Ultrasonic Spatial Target Localization Using Artificial Pinnae of Brown Long-eared Bat

**DOI:** 10.1101/2020.03.02.972802

**Authors:** Sen Zhang, Xin Ma, Zheng Dong, Weidong Zhou

**Author notes:** Corresponding author: Xin Ma.

## Abstract

Echolocating bats locate a target by echolocation and their performance is related to the shape of the binaural conformation in bats. In this study, we developed an artificial sonar system based on the vertical sound localization characteristics of the brown long-eared bat (*Plecotus auritus*). First, using the finite element method, we found that the beam of the first side lobe formed by a pinna constructed according to that in the brown long-eared bat shifted in an almost linear manner in the vertical direction as the frequency changed from 30 kHz to 60 kHz. We established a model of the relationship between the time-frequency features of the echo emitted by brown long-eared bats and the spatial direction by using the pre-trained neural network. We also developed a majority vote-based method called sliding window cumulative peak estimation (SWCPE) to optimize the outputs from the neural network. In addition, an L-shaped pinna structure was designed to simultaneously estimate the azimuth and elevation. Our field experiments indicated that the binaural conformation and relative binaural orientation both played vital roles in spatial target localization by these bats. Accurate echolocation can be achieved using a simple binaural sonar device even without binaural time difference information.

## I. Introduction

The disruptive development of mobile and flying robots such as automatic guided vehicles and unmanned aerial vehicles demands new sensory approaches for target search and obstacle avoidance. Machine vision-based methods can satisfy the basic requirements in favorable lighting environments but variables such as darkness, fog, and smoke hinder their wider application. Sonar sensing can effectively complement the commonly used vision sensing techniques in robot applications. In traditional sonar systems, transmitter and receiver arrays are used widely for navigation and localization, but the excessive number of sensors included in these systems often causes difficulties during installation and layout. In contrast, the bat echolocation system is an extremely compact sonar system with only two sound receivers and one speaker, which would obviously be preferable for robot applications because of its simplicity, low cost, and high performance [1] [2].

The excellent navigation and target localization capabilities of bat sonar have attracted much attention from researchers who have focused on various aspects. In particular, the relationships between echolocation and the auditory neurons or auditory cortex [3] [4] [5], have been investigated in order to identify a suitable echo processing method for navigation and localization by observing the activities of the auditory nerve and auditory cortex during echolocation by bats. In addition, studies of the behavior of bats [6] [7] [8] [9] [10], have provided great insights into the intrinsic mechanism responsible for navigation and localization by bats. Moreover, investigations have considered the acoustic roles of the physiological structures of bats, including the sound field characteristics related to the facial physiological structure [11], auricle structure [12], and vocal tract [13]. All of these studies have above provided important insights to facilitate the artificial reproduction of echolocation by bats.

Some interesting results in bat research have been physically reproduced and even implemented in practical applications. Most two-dimensional target localization methods based on binaural systems have been implemented based on the interaural time difference (ITD) [14] [15] or interaural intensity difference (IID) [16], whereas the majority of the three-dimensional (3D) target localization systems have employed multi-receiver designs [17] [18]. However, these types of applications are not fundamentally different from the conventional sensor array methods. Some reproduction techniques exploit the unique sound field characteristics determined by the physiological structure of bat ears, which more strictly imitate the localization effect of bat sonar. For example, Chiu et al. conducted experiments based on the behavior of big brown bats (*Eptesicus fuscus*) and found that the tragus is related to vertical sound localization [19]; Kuc designed an experimental model based on acoustic mirror formed by rotating a lancet, which exhibited an elevation versus notch frequency sensitivity [20];and Schillebeeckx et al. [21] used two microphones and one ultrasonic emitter, conducted the experiments of echolocation for a spatial target in the laboratory. In our previous study [22],using a finite element method (FEM) [23], We found in a specific wide range, the time-frequency characteristics of sonar in long-eared bats had a strong relationship with the vertical direction (i.e., elevation angle). In the present study, we estimated the angle of the echo using another approach inspired by the sound field characteristics of the pinnae in brown long-eared bats based on FEM simulations. The novelty of this work with respect to previous realizations lies in the utilization of the distinguishing ability of the brown long-eared bats in vertical space and its simplicity in real-world application. This study obtained the following three main conclusions. The rest of the paper is structured as follows. Section II describes the methods used in our work. Section III describes the results of our experiments, including the results of simulation and field experiments. Section IV discusses the method in our experiments for target echolocation, and Section V draws a conclusion on the system and its potentiality of application.

## Materials and methods

### A. Numerical analyses of *Plecotus auritus*

In order to obtain more evidence to inspire the design of an accurate artificial bat-like sonar system, we conducted numerical simulations of the pinnae in *Plecotus auritus* by using a FEM and Kirchhoff integral [24] to analyze the spatial frequency characteristics. In particular, 3D digital models of *Plecotus auritus* pinnae were obtained by digital image processing [25] using computed tomography scans of pinnae (Fig 1B). We placed a sound source in the inner ear canal in the numerical model and performed numerical calculations based on the FEM. First, we used FEM and Kirchhoff integral to obtain the beam pattern for the digital ear in the simulation. The acoustic near field inside a cuboid-shaped volume surrounding the ear was then calculated using a FEM comprising linear cubic elements derived directly from the voxel shape representation. Finally, the far-field directivity pattern was calculated by projecting the complex wave field amplitudes outward onto the surface of the computational domain of the FEM using a Kirchhoff integral formulation.

**Fig. 1.**
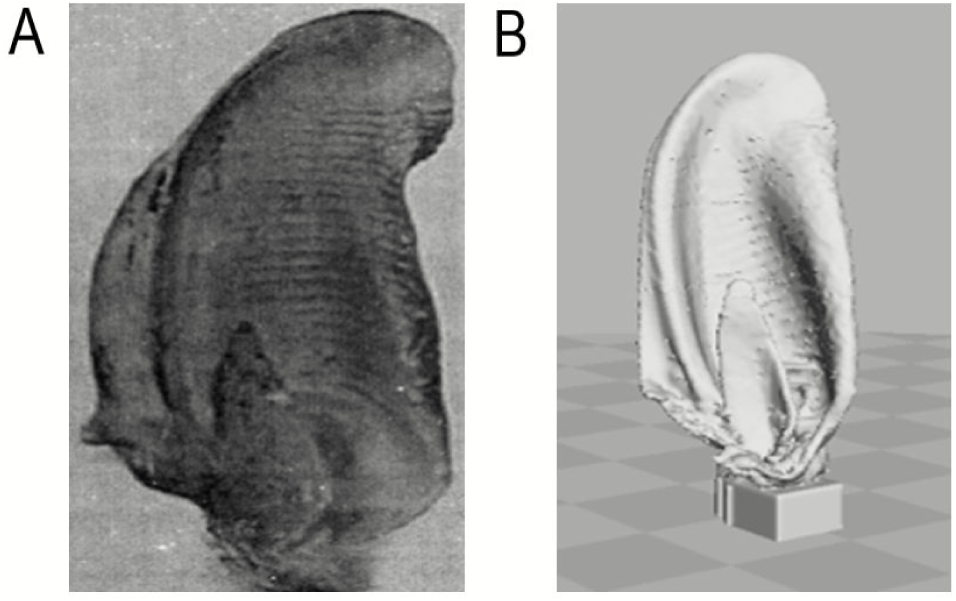
Pinna of brown long-eared bat. (A) Photo of the pinna of brown long-eared bat. (B) Three-dimensional model of brown long-eared bat pinna.

### B. Artificial bat-like Device

The artificial bat-like ears were produced using a 3D printer according to a 3D digital model obtained based on pinnae samples taken from the carcass of a brown long-eared bat. To avoid damage during assembly and to allow the fixation of a microphone, the size of each printed artificial bat-like ear was three times larger than the original ear. The frequency range of *Plecotus auritus* ears is 2060 kHz, so the frequency range used in the experiments was adjusted to 5kHz to 20 kHz according to the scale model principle [26]. A pair of ultrasonic microphones (SPU0410LR5H-QB, Knowles Electronics, Itasca, Illinois, USA) were placed inside the pair of artificial ears and insulating glue was smeared in the gaps between the microphones and the pinnae to prevent outside sound waves entering the microphones from the bottom of the model. The artificial ears were then fixed to a rotating platform and tilted forward 40°(Fig 2E). A stepping motor was mounted under the rotating platform to facilitate rotation of the ears and to measure positioning information at the azimuth. An ultrasonic loudspeaker (UltraSound Gate Player BL Light; Avisoft Bioacoustics e.K., Glienicke, Germany) was fixed under the stepping motor (Fig 2A,B).The target was fixed on the green spots and the device was located in the eight different locations in turn to obtain the training data. Yellow spots A, B, and C were used for locating the device to collect the testing data. The layouts of the target and device in the training and testing processes are illustrated in Fig 2F.

**Fig. 2.**
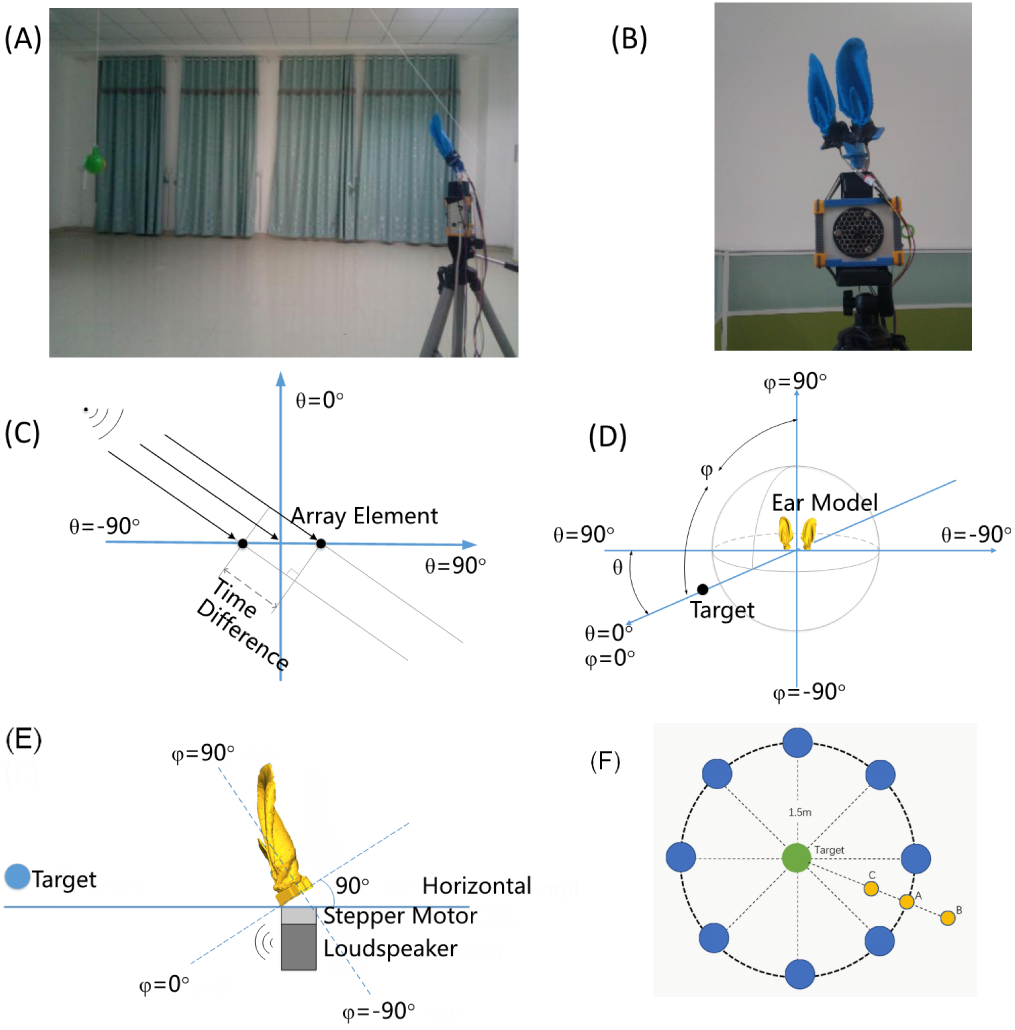
Bat-like sonar and layout used in the single target localization experiments. (A, B) Target, parallel erect pinnae device, and the environment. (C) Top view of platform. (D) Spherical coordinate. (E) Side view of the artificial pinnae.(F) Top view of the target and device during training and testing.

### C. Data acquisition

All of the data acquisition experiments were conducted in an experimental chamber (8 m (length) 6 m (width) 3.6m (height)) (Fig 2A). No sound insulation was present in the chamber. The acquisition parameters are listed in Table I. The target for measurement was a small ball made of rubber with a diameter of 11cm and it was suspended by a string. The frequency response of the ear model in different directions could be measured by controlling the height of the target and the rotation angle of the motor (Fig 2A,2C,2D). The ultrasonic signal acquisition and processing device was a signal acquisition card (PXIe-6358 and PXIe-1082; National Instruments, Austin, Texas, USA; sampling at 100 KS/s), which allowed multi-channel synchronous signal acquisition. The signal emitted by the brown long-eared bat is a type of linear frequency-modulated pulse signal, which has similar frequency characteristics to a chirp [27], so we set the ultrasonic loudspeaker to emit a chirp pulse signal (see Fig 3A) in our active target echolocation experiments in order to imitate the pulse emitted by the brown long-eared bat, and its time-frequency intensity is shown in Fig 3D. We set the signal acquisition card in the work mode for two-channel synchronous signal acquisition in order to collect the binaural signals in a synchronous manner. Details of the signals emitted in the active echolocation experiment are also listed in Table I.

**TABLE I.** Experimental parameters.

| ID | Name | Comment |
| --- | --- | --- |
| 1 | Sampling rate | 100 kHz |
| 2 | Model size of bat ear | Three times the original size |
| 3 | Microphone distance | 5 cm |
| 4 | Transmission signal | Linear frequency modulation signal |
| 5 | Frequency range | 5-20kHz |
| 6 | Active signal duration | 5 ms |
| 7 | Alteration mode of azimuth | By altering the height of the target (small ball). |
| 8 | Alteration mode of elevation | By rotating the pinna. |
| 9 | Scope of azimuth alteration | -90° to 90° (step length: 10°) for estimating the elevation in parallel erect pinnae experiments; -28° to 28° (step length: 7°) for the orthogonal pinnae experiments. |
| 10 | Scope of elevation measurement | 20 to 55° (step length: 5°) in parallel erect pinnae experiments, 1268° (step length: 7°) for the orthogonal pinnae experiments. |

**Fig. 3.**
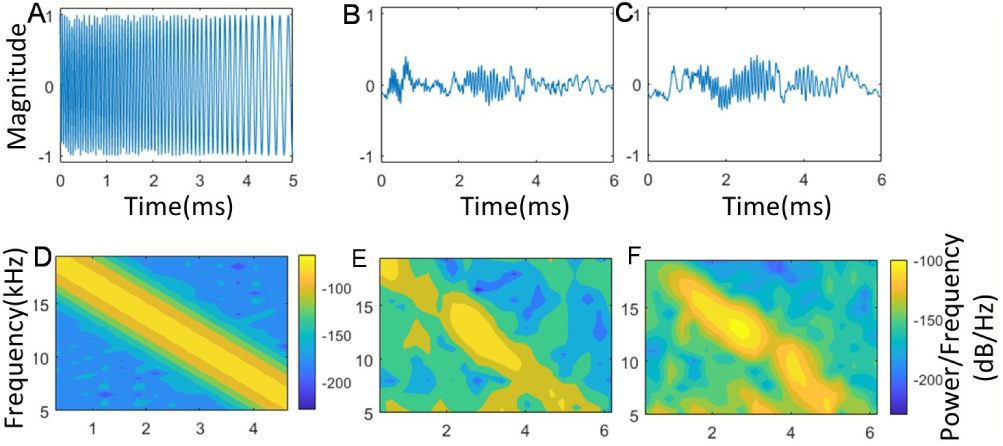
Example of a signal emitted by the loudspeaker and echoes received by the two ears. (A,B,C) Emitted signal and echoes received by left and right microphones. (C)Top view of the platform. (D,E,F) Spectrograms for the emitted signal and echoes received by the left and right microphones.

### D. Feature extraction

The signals received by the left and right microphones are shown in Fig 3B and 3C. Zero crossing rate endpoint detection with center clipping was conducted to detect the beginning position of the echo. The effective signals were then transferred into time-frequency representations as:

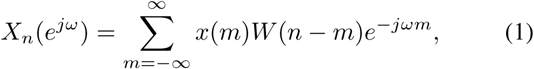

where *x*(*n*) is the signal received after the endpoint detection process and *W* (*n*) is the window function, which shifts the sound signal by a step length on the time axis. We used Hamming windows with a length of 1 ms (100 samples) as the window function and the shift step was half of the window length. The time frequencies at 0.5 kHz frequency resolution for the pulse signals received by the left and right microphones are shown in Fig 3E and 3F.

The spectrogram energy in the emitted pulse signal was mainly concentrated near the diagonal line, so the spectrogram energy for the echo was also concentrated near the diagonal line. Thus, we set a threshold to suppress the effects of interference components while maintaining the value constant on or near the diagonal line in the spectrogram. The spectrogram was then restricted to one vector by summing the values along the frequency channels. After the treatment, a time-frequency representation with length T was transformed into a vector with *n* elements, as described in Equation 2. In our experiment, we selected *n* = 30 and connected the vectors from the left and right echo signals to obtain one vector with 60 elements as the input for the classifier.

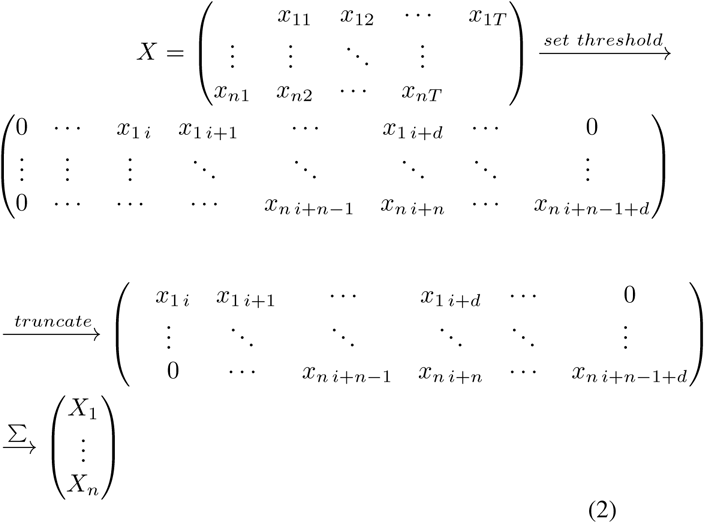

### E. Neural network for estimating the angle of the spatial target

Artificial neural networks are used widely in pattern recognition tasks as efficient methods. At present, deep learning is very popular because of its good classification performance, but we selected the traditional back-propagation (BP) feed-forward neural network as an estimation tool because of its simplicity and practicality, but also because of its convenient regression function. We applied the BP feed-forward neural network to three tasks comprising elevation estimation in the case of parallel erect pinnae, elevation in the case of orthogonal pinnae, and azimuth estimation. The BP feed-forward neural networks used in the three tasks had the same structure (Fig 4A), which comprised an input layer with 60 neurons (30 + 30, i.e., the features extracted from the echo signals for the left and right artificial ears where fed directly into the network), a hidden layer with nine neurons, and an output layer with one output neuron. The three layers were fully connected in the BP neural network. Tan-sigmoid and pure linear were selected as the activation functions in the hidden layer and output layer, respectively. In the training phase, the neural network learned the characteristics of the time-frequency patterns reflected by the target at different angles (elevations or azimuths). The network was trained using the LevenbergMarquardt optimization algorithm. After training, the time-frequency patterns generated from untrained ultrasonic echoes were fed into the network. The structure of the neural networks used in the three tasks was the same but the normalization functions were different because the output angles differed. The three types of outputs were linearly transformed into activities between 0.05 and 0.95, and they are shown in Fig 4B, 4C, and 4D, respectively. In the estimation process, the output neuron activities were linearly re-transformed according to the activity functions and they were used as the angle estimates for the respective inputs in each task.

**Fig. 4.**
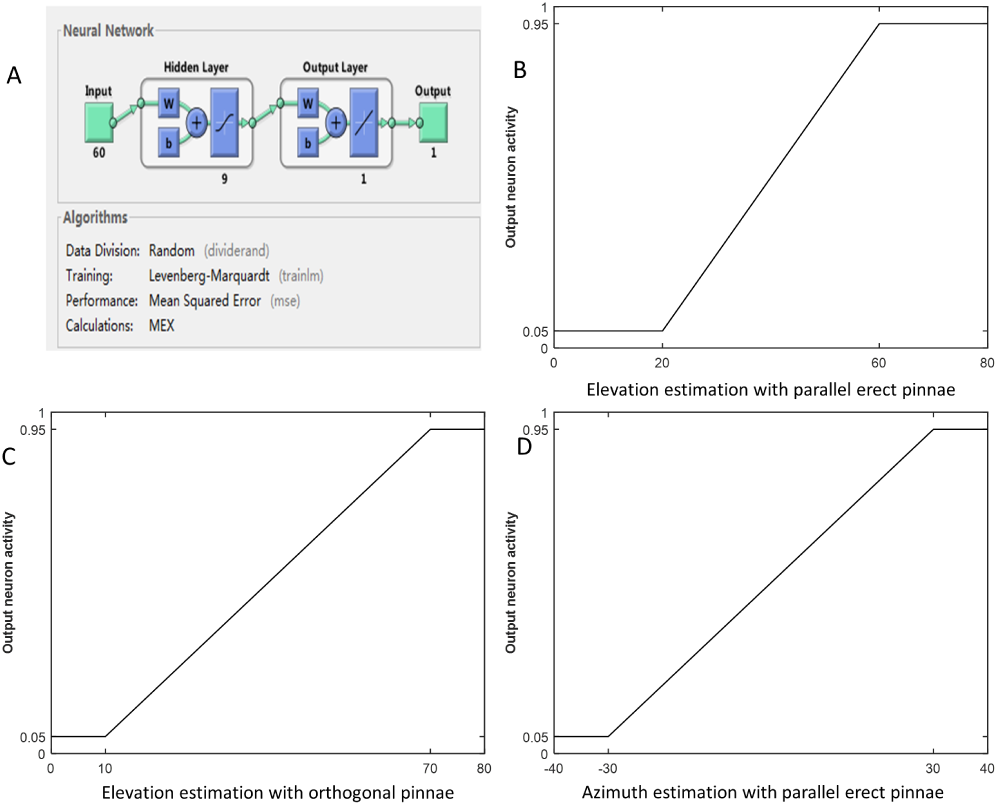
Back-propagation feed-forward neural network used in our system. (A) Structure of the neural network. (B), (C) and (D) Transformation functions between the angle value and output neuron activity for training and testing in different tasks.

### F. Sliding window cumulative peak estimation (SWCPE) under multi-pulse input

The activities of the output neurons indicated the values of the azimuth or elevation during the analysis of one pulse. If multiple pulses were analyzed for one target by the same classifier, multiple estimated values were exported. In general, further optimization could be conducted based on the multiple estimated values and the optimized values obtained by the method had a higher level of confidence. This principle may explain why bats locate targets based on a pulse train. In order to imitate the actual signals that are generally emitted by bats in the form of a pulse train, we developed a novel direction angle estimation enhancement method called SWCPE for obtaining near-optimal estimates based on multiple relatively rough estimation values. For *n* initial estimate values, the P-levels SWCPE process is explained as follows.

1. For *n* estimated values (each value represented one sample) obtained by a certain method, such as a BP neural network, the first level moving window with length L moved over the range of all *n* samples with a step length l. The starting position of the window (left edge) was aligned with the point with the minimum value and the end position (right edge) was the sample point with the maximum value. Each point in the range was set with an initial token value *y* of 0.
2. When the window moved one step, the y value for each point in the window increased by the number of samples in the current window. After the window reached the end, we calculated the point with the maximum *y* value as the optimal value of the first-order moving window. If more than one point had the maximum value, we selected the intermediate point between the left and right points with the maximum value as the optimum solution.
3. In general, the first-order moving window was used to obtain an optimized scope that could include the optimum solution with the maximum probability. Next, *L/*2 length second-level window and *L/*4 length third-level window estimation were performed up to a p-level window estimation to obtain a narrower search scope. The number of levels determined the accuracy of the estimate. In principle, the accuracy of the estimate was less than the half of the windows width.

## Results

### Numerical analysis results: Spatial acoustic characteristics of digital brown long-eared bat pinnae

Frequencies spanning the entire frequency range (from 20 kHz to 60 kHz with a step size of 1 kHz) known to be covered by the first harmonic [28] of the biosonar pulses were analyzed using the numerical method (Method A). The first side lobe in the beam pattern (Fig 5A, 6B) exhibited a frequency-driven scanning characteristic and its power was relatively strong when the frequency exceeded 32 kHz. When the frequency was less than 30 kHz (Fig 5C), the half-power beam width curve was relatively large whereas the power of the side beam was low and the orientability of the lobe was not concentrated, which resulted in low directional resolution. The beam direction of the first side lobe shifted along the elevation in an almost linear manner as the frequency changed from 30 kHz to 60 kHz, whereas the azimuth of the side lobe remained almost stable (Fig 5D). The peak of the beam for different frequencies appeared to alternate along the elevation, thereby suggesting that in this frequency band, the combination mode of the frequency intensities changed greatly in the direction of elevation, and this implied that the vertical resolution was strong under this frequency combination with a potential functional relationship.

**Fig. 5.**
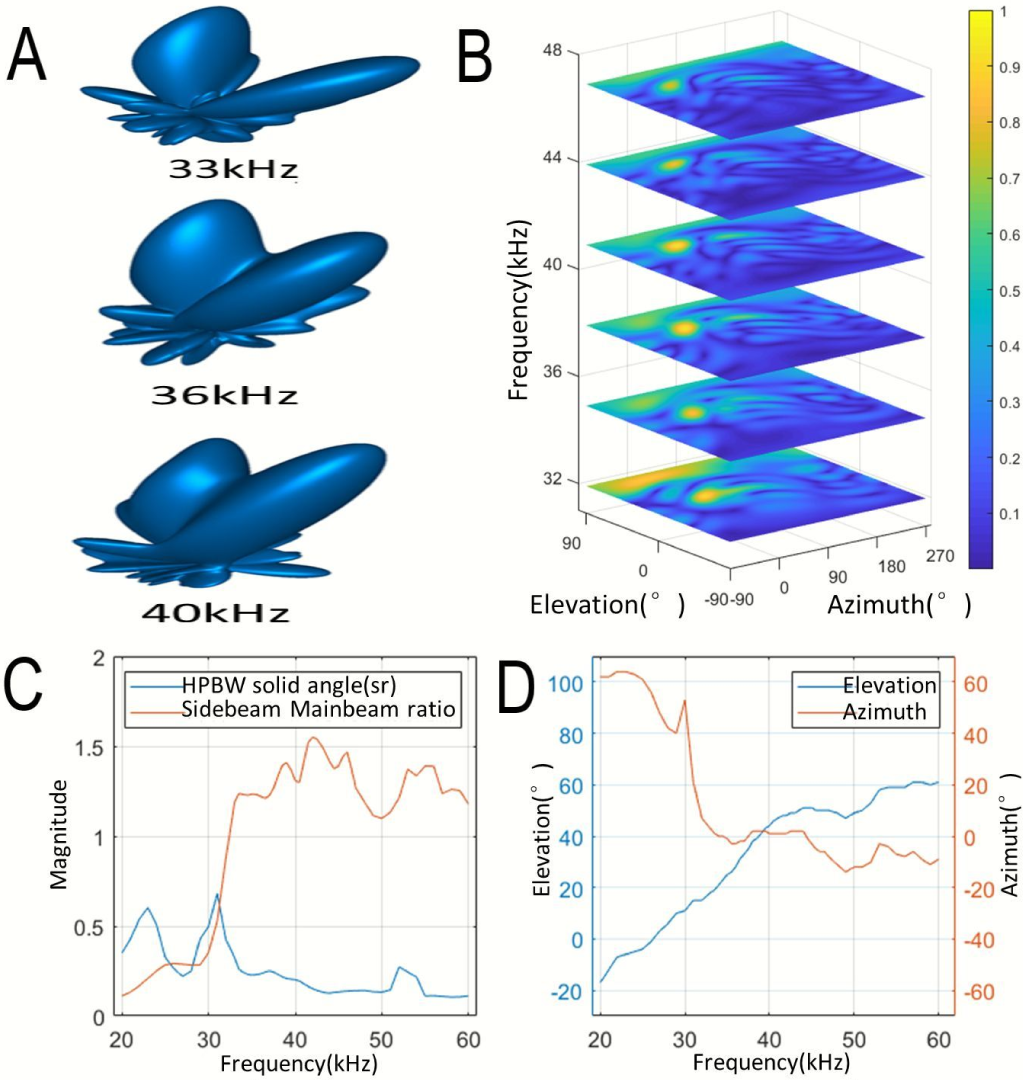
Acoustic characteristics of the digital pinnae. (A) Beam pattern of the pinna. (B) Beam pattern of the sound field under different frequency. (C) Half-power beam width and energy ratio for the side lobe and main lobe. (D)Elevation and azimuth for the side lobe with different frequencies.

**Fig. 6.**
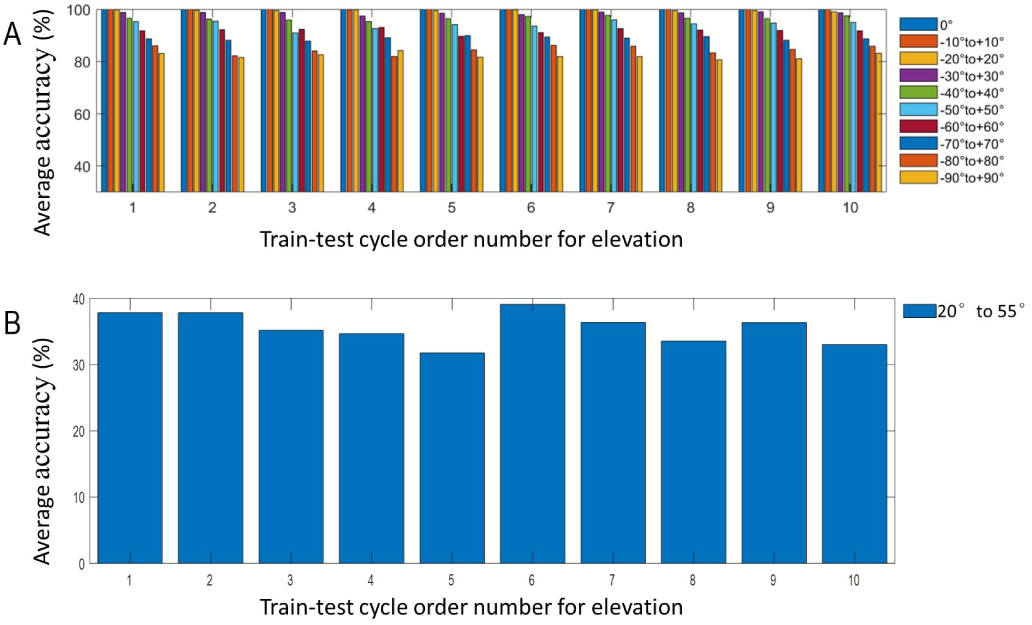
Single pulse estimation results obtained with parallel erect pinnae. (A) Average accuracy (under ± 5° error) of the elevation estimates in 10 cross-validation cycles. The 10 histograms for each cycle represent the limited scopes of the azimuth for the training set ranging from ± 0° to ± 90°.The vertical axis represents the average precision ratio where the value with an error less than 5° was considered to be the correct result.(B) Average accuracy (under ±5° error) of the azimuth estimations in 10 cross-validation cycles. Only one histogram in each cycle represents the result obtained under the limited scope of 20° to 55° for the elevation.

### Artificial bat-like sonar device experiment: Vertical spatial resolution of the frequency combination and application to target localization

A potential relationship between the frequency combination and spatial direction was demonstrated by the simulations, but we had little information about how wide this relationship might spread in the hemispherical space and how to apply it in the physical space. Thus, physical verification was still required for the spatial resolution of the frequency combination in the vertical direction and the range of the width also had to be clarified. Thus, a bat-like device was designed to estimate the angle of the spatial point-like target (Fig 2A) by mimicking the function of the ears of the brown long-eared bat in order to clarify whether the consistency width range of the frequency combination changed with the vertical direction. The detailed experimental parameters are shown in Table I.

#### A. Target elevation estimation using parallel erect pinnae under different width ranges

We used the artificial bat-like sonar device to emit and receive acoustic signals with the two artificial pinnae pointing upward and collected data using the data acquisition device (Method C), as summarized in Table I. Inspired by the high resolution of the pinna and the potential functional relationship implied by FEM analysis, a BP neural network was employed to estimate the continuous position in the vertical direction. The detailed structure of the BP neural network used in our study is presented in Method E.

##### 1) Single pulse target elevation estimation under different width ranges

The simulation results demonstrated the differentiation of the frequency combination with the elevation in a linear region. To test whether the properties of this frequency combination were related to the elevation in a wider range, we combined 10 data sets of spatial feature samples according to the different widths in the horizontal direction, and estimated the elevations in each data set. Single pulse and different sets of data categories under different azimuth scopes for every statistic (Fig 6A) were used to evaluate the accuracy of the elevation estimated under different width ranges. In order to obtain reliable results, the sonar device was located at eight different sites (blue circles in Fig 2F), where the small ball used as the target was located at the center, as indicated by the green circle in Fig 2F. The horizontal distance between the ball and the sonar device was 1.5m. The training data and testing data were different but they came from the same data set, as explained in Method G. The values of − *N*° to +*N*° represented the scope of the limited azimuth angle from the left and right (Fig 6A). Tenfold cross validation was conducted to obtain reliable results for each − *N*° to +*N*° scope. The amounts of data in each training and testing set were related to this range, i.e., (((*N/*10) × 2 + 1) × 8) × 0.9 for training and (((*N/*10) ×2 + 1) × 8) × 0.1 for testing. In order to ensure that the distribution of the data was reasonable, the training data and test data were evenly sampled from eight different locations. The mean results obtained by tenfold cross-validation are shown in Fig 6A. The average precision was nearly 100% with the scope from − 20° to +20° of the limited azimuth. The average accuracy exceeded 90% when the limit angle of the azimuth was − 60° to +60° and the average accuracy exceeded 80% even when the limit angle of the azimuth was − 90° to +90°. These findings illustrated the good wide-angle effect on elevation estimation and the high accuracy in the middle direction of the azimuth, which are essential for target detection. In contrast, the average accuracy of the azimuth estimation was only about 35 percent when the limited angle of elevation was 20° to 55° (Fig 6B). Thus, the azimuth estimates were clearly not as good as the elevation results, thereby indicating that the changes in the features with the azimuth lacked regularity compared with the elevation. The experimental results confirmed the results obtained in the simulation experiment and they clarified the effective width range in the vertical direction.

##### 2) Improving the echolocation accuracy using a pulse train

Many studies have shown that bats can enhance their echolocation ability by transmitting a pulse train [29] [30] [31] [32] [33]. Thus, we used a pulse train estimation method to imitate this behavior and the SWCPE method (Method F) was developed to obtain accurate results. Increasing the pulse number in each train could effectively compensate for the reduced accuracy of the elevation estimation caused by the increase in the azimuth range. When the number of pulses in one pulse train reached 20, the elevation estimation accuracy was more than 95% under an error standard of ± 3° (middle panel in Fig 7B) within an azimuth range of ± 50° and more than 90% under an error standard of ± 5° within a large azimuth range up to ± 90° (bottom panel in Fig 7B). Thus, by using the pulse train, the bats could obtain higher accuracy and a better wide-angle range for target localization.

**Fig. 7.**
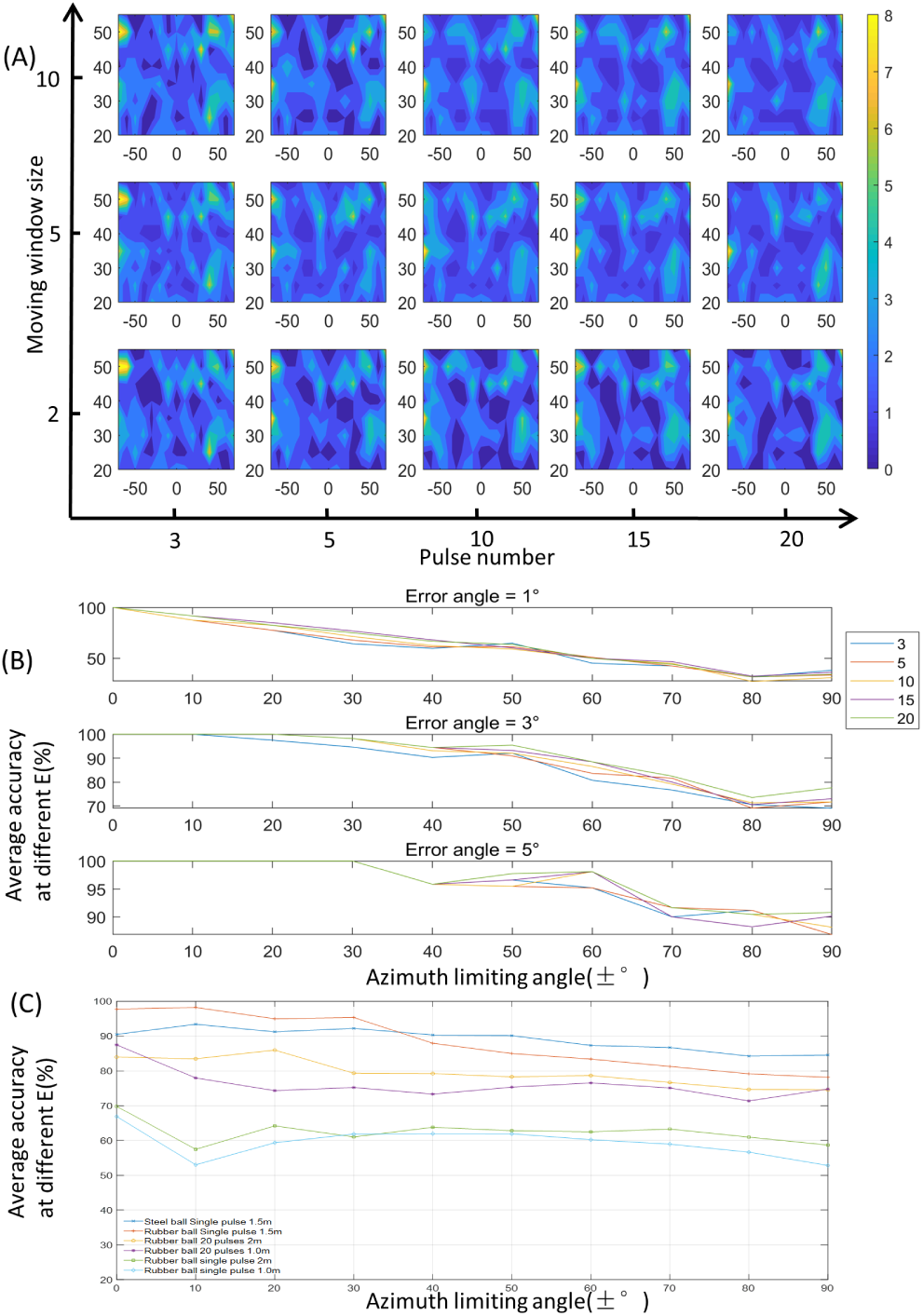
Pulse train estimation results obtained with parallel erect pinnae under different conditions. (A) Spatial distribution of the elevation estimated using a pulse train under conditions with different numbers of pulse trains (from left to right, the number of pulses in a single pulse train range from three to 20) and different sliding window levels (from top to the bottom, the numbers of levels in SWCPE are one, two, and three, and the widths of the sliding window at each level are 10, five, and two, respectively; see Method F). The horizontal and vertical axes in each small colormap represent the limited azimuth and estimated elevation (range from 20° to 55°), respectively. (B) Average elevation estimation accuracy using pulse trains under different error criteria (from top to bottom: ± 1°, ± 3°, and ± 5°, respectively) and under different limited azimuth ranges. The different colors represent various numbers of pulses in a single pulse train. (C) Average accuracy of elevation estimation for a steel ball with a single pulse at 1.5 m, rubber ball with a single pulse at 1.5 m, rubber ball with 20 pulses at 2 m, rubber ball with 20 pulses at 1.0 m, rubber ball with a single pulse at 2 m, and rubber ball with a single pulse at 1.0 m.

##### 3) Generalization tests

Robustness testing methods were designed to verify the generalizability of the system used for estimation. The training data were the same as those used for single pulse target elevation estimation. The training process was also similar to the process employed for single pulse target elevation estimation, except no data in the training set were retained for use as test data. Thus, for every limited scope of the azimuth, the amount of data used in each training set was (((*N/*10) × 2 + 1) × 8). The generalization testing samples were collected in the same experimental chamber but under conditions where the artificial bat-like devices were placed outside the eight sites used in the training process (e.g., sites A and B in Fig 2F). The neural network used for the test was trained with the 90° limited scopes of the azimuth and under the 1.5-m condition, as described above. Cross-validation was not conducted in the testing process. SWCPE (Method F) was also used for multiple pulse testing to obtain the final estimated values. Fig 7C shows the performance under six conditions. The results were similar for the steel ball used as a reflector in this experiment, although it was not used for training. The results were similar although the steel ball was not used for training. The performance was greatly influenced by the distance (the green line with a cube and blue line with a diamond). The intensity of the echo was larger at 1 m than 2 m, but the performance was not better than that with 2 m. The multiple pulse chains effectively improved the performance with distances of 1 m and 2 m, and the improvements were slightly greater with 2 m than 1 m.

#### B. Target angle estimation using orthogonal pinnae

In the elevation estimation process described above, the two pinnae of the brown long-eared bat were parallel to each other. We observed that the two pinnae of the brown long-eared bat often stretch to a certain angle when hunting for prey. If the angle is 90°, the orthogonality of the two pinnae can be used to obtain the aspect angles in the two orthogonal directions. The orthogonal pinnae of the artificial brown long-eared bats used for target localization are shown in Fig 8. Except for the differences shown in Table I, the other measurement conditions were the same as those employed in the generalization tests.

**Fig. 8.**
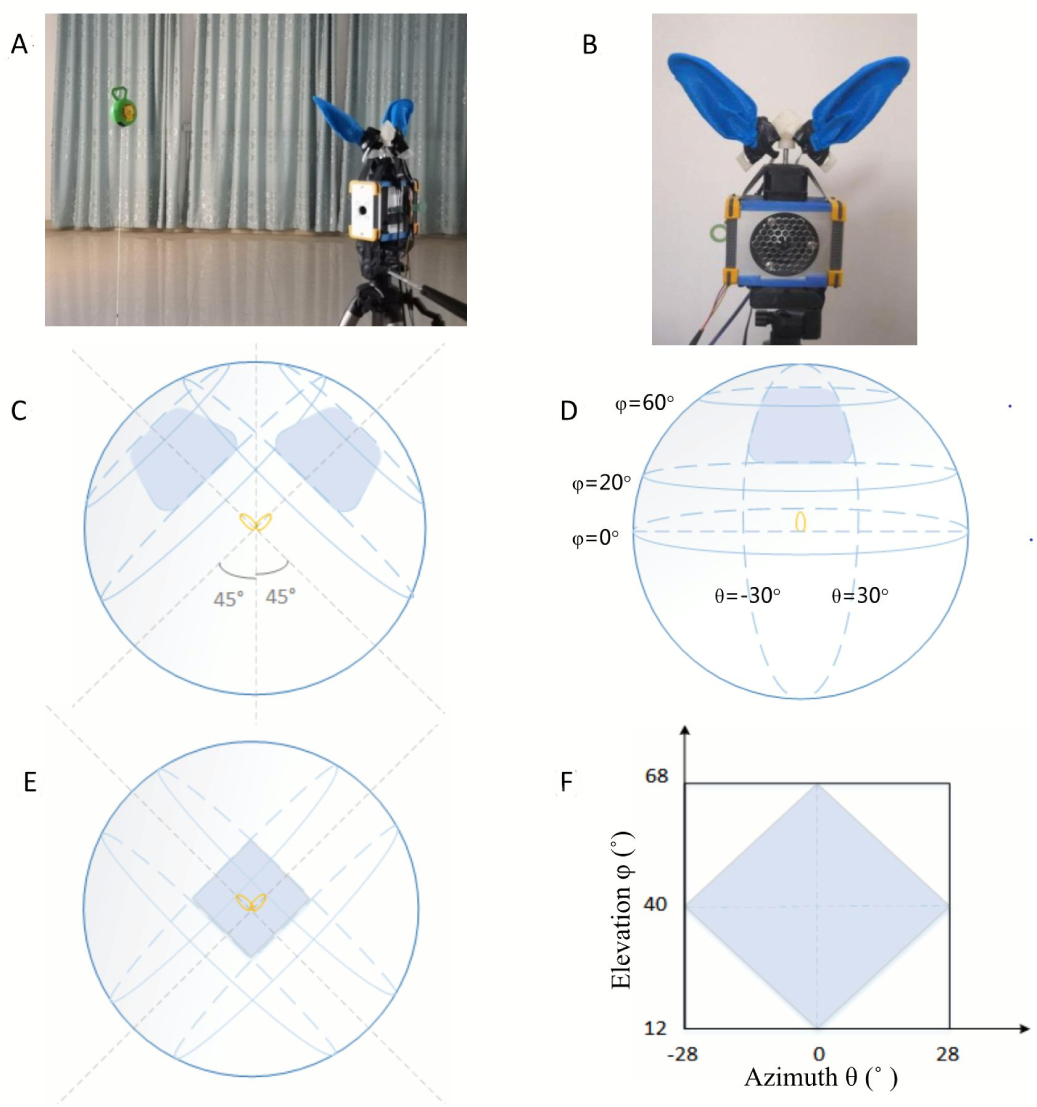
Orthogonal pinnae active sonar device and design of the experiment. (A, B) Images of the target, orthogonal pinnae device, and environment. (C, D, E, F) Process employed to form the effective target region. Before tilting the pinna, the effective ranges of the beam patterns for the two pinnae corresponded to the shaded regions in panel C. The estimated elevation regions for each pinna are shown as the shaded regions in panel D. Each pinna tilted forward 40° to form the overlapping region, which is shown as the shaded region in panel E. The actual statistical scope was − 28° to +28° in the azimuth and 12° to 68° in the elevation.

The results in Fig 9 demonstrated that elevation estimation and the azimuth estimation could be performed simultaneously with the orthogonal pinnae, thereby indicating the effectiveness of the proposed method. In the box plots in Fig 9A and 10B, the red horizontal lines denote the median estimated angles, and these result show that 50% of the estimated angles were located in the blue box, and thus both the elevation and azimuth estimation method achieved good results. The standard deviations of both the elevation and azimuth results remained fairly stable at various angles. This result indicated that although the signal-to-noise ratio (SNR) was poor at high elevations or in the marginal azimuth, the correlations between the sweeping frequency and elevation were strongly maintained. Similar to the case with parallel erect pinnae, the pulse train estimation method was used to correct the deviations in the estimated angle and to enhance the accuracy. Fig 9C shows the localization results under different pulse trains. For all of the pulse trains, the percentages increased as the number of pulses increased. For the pulse train with 10 pulses, more than 60% of all the target angles were estimated with an error of 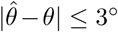 and 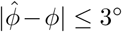, more than 91% with an error of 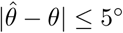 and 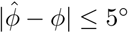, and the error rate was only less than 1% when the demand was 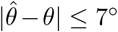; and 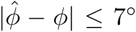. Clearly, using a pulse train improved the estimation accuracy. Fig 9D shows the performance under 2.0 m using the training data obtained with 1.5 m. The different distance decreased the match between the training data and test data, but the pulse chain effectively compensated for the loss of the performance in the case of orthogonal pinnae.

**Fig. 9.**
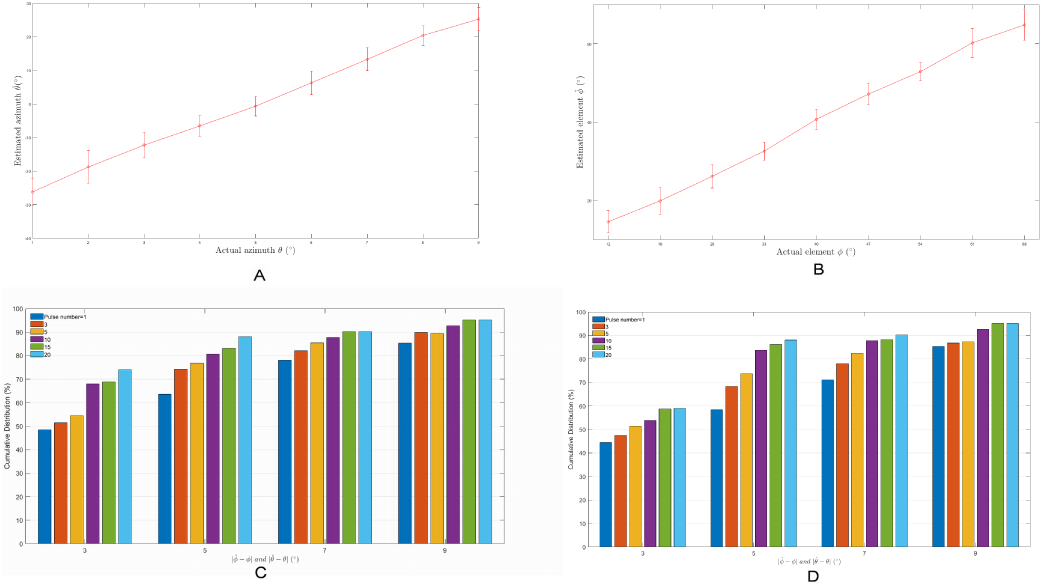
Elevation and azimuth estimated with the active orthogonal pinnae. (A, B) Comparisons of the azimuth and elevation estimates in the single pulse test with the actual values. (C) Cumulative distributions for the corresponding azimuth and elevation errors. The abscissa axis represents the errors for both the azimuth and elevation to meet. (C) Cumulative distributions for the corresponding azimuth and elevation errors with 1.5 m. (D) Cumulative distributions for the corresponding azimuth and elevation errors with 2.0 m.

## Discussion

In this study, we developed a strictly bat-like sonar system inspired by the brown long-eared bat (*Plecotus auritus*). The sonar system comprised one emitter and two microphones fitted with artificial pinnae to imitate the physical conformation of the sonar system in bats, and the signals emitted were similar to those produced by brown long-eared bats. In contrast to other bat-like sonar systems, our strictly bat-like sonar had almost the same pinna construction, while the same sound was emitted and received as the prototype. This strict bat-like method for reproducing the behavior of bats can provide further insights into the true sonar mechanism employed by bats as well as serving as a validation method.

Schillebeeckx et al. established a set of artificial pinna models for 3D target location [34] as an important advance in the development of strictly bat-like sonar. They used a Bayesian classification method to separately model all of the positions in hemispheric space at a resolution of 1°. Their experiments demonstrated the feasibility of a strict bat-like model for spatial localization. The process determined in their experiments suggested that the echo transfer functions in each direction are independent of each other, i.e., the input features need to be calculated in every model. We suggest that this method is theoretically suitable to some degree for all types of bionic models of pinna, including the human ear. However, in practice, the results obtained by this method for each type of auricle will be quite different, because the differences in the feature distributions in different directions are not always conspicuous, and thus the model parameters will be too close in different directions.

The main difference between our method and that employed by Schillebeeckx et al. is that we considered the characteristics of the long-eared brown bat where the spatial sound field beam clearly changed with the frequency in the vertical direction (central axis of the auricle). This effect was clearly demonstrated in our FEM simulations. However, the scope of the width of the effect and the effect on the horizon were not clear based on the results obtained in our FEM simulations. Experiments using our artificial bionic ear model showed that the changes in the echo features in the vertical direction were related to the combination of the frequency components and this vertical correlation was maintained in a wide range up to ± 60° (Fig 7B). The correlation was still present in the horizontal direction (Fig 6B), but it was significantly weaker than that in the vertical direction.

We also identified the intrinsic relationship between the time-frequency features of the echoes used by the brown long-eared bat and the spatial direction. Spatial angle information could be obtained from the echo, and thus the echo must contain angle information. Thus, the echo features were directly related to the angle information. According to the relationship between the features of the echolocation-related transfer function (ERTF) and spatial position, if we use *X* to denote the ERTF features and *f* to express their distribution in two-dimensional space (*θ, φ*), then *X* can be written as follows.

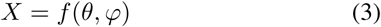

Our experiments suggested that in a certain range of azimuth angles, a function *g*(*X*) exists that is only related to *φ*:

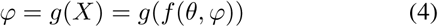

where *θ* and *φ* denote the azimuth and elevation of a target, respectively. Thus, it is necessary to determine the function in terms of the relationship between the angle information and features in the orthogonal coordinate system. This function is usually difficult to obtain, but the relationship can be approximated by the BP-based feed forward neural network as:

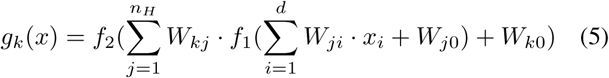

where *k* is the number of output variables, and *f* 1 and *f* 2 are discrimination functions in the hidden and output layers, respectively. In the case of parallel erect pinnae, we can obtain:

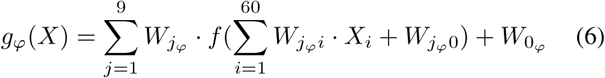

where *f* is the discrimination function in the hidden layer:

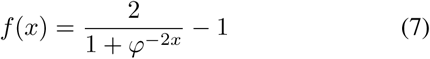

For orthogonal pinnae, we can obtain *g*_*φ*_(*X*) and *g*_*θ*_(*X*) separately using two independent neural networks.

Feature distortion had important effects on the estimated results and a change in the SNR ratio was one of the factors responsible for feature distortion, as also mentioned by Schillebeeckx et al [35]. However, for the results estimated directly for the targets made of different materials and at different distances (Fig 8C), the SNR ratio was not the only factor responsible for distortion. In particular, the SNR ratio for the echo from the steel ball was much stronger than that for the wave reflected from the rubber ball, but there was no significant difference in the estimates for the steel ball and rubber ball. When we tested the targets located at 2 m using the network trained at 1.5 m, the results were poor even though the SNR of the reflected echo at 1 m was better than that for the target at 2 m. No other form of distance compensation was conducted in our experiments apart from using a simple classification decision method for multiple outputs. However, a real bat may have the ability to compensate for distance. Effectively compensating for the loss of echo features at different distance should be addressed in future research.

## Conclusion

In contrast to other spatial angle classifier estimation methods, our proposed algorithm only used a one-dimensional label, i.e., either the elevation or azimuth, as the output from our classifier model. Our experiments also suggested that the proposed structure and model are suitable for accurate spatial target echolocation with a limited computational load in a realtime system.

## Acknowledgments

This work was supported by the Shenzhen Science and Technology Research and Development Funds (Grants No. JCYJ20170818104011781) and the Key Research and Development Program of Shandong Province (Grants No. 2017GGX10113 and 2019GGX101063). We also thank all the anonymous reviewers for their helpful and stimulating comments on earlier versions of the manuscript.

